# Monte Carlo track-structure simulation of the impact of Ultra-Hight Dose Rate and oxygen concentration on the Fenton reaction

**DOI:** 10.1101/2025.05.13.652705

**Authors:** M. Chaoui, Y. Tayalati, O. Bouhali, J. Ramos-Méndez

## Abstract

**Background:** Preclinical investigations studies have shown that FLASH radiotherapy (FLASH-RT), delivering radiation in ultra-high dose rates (UHDR), preserves healthy tissue and reduces toxicity, all while maintaining an effective tumor response compared to conventional radiotherapy (CONV-RT), the combined biological benefit was termed as “FLASH effect”. However, the mechanisms responsible for this effect remain unclear. Research demonstrated that oxygen concentration contributes to the FLASH effect, and it has been hypothesized that Fenton reaction might play a role in the “FLASH effect”.

**Purpose:** We propose to investigate the effect of ultra-high dose rate (UHDR), compared to conventional dose rates (CONV), on the Fenton reaction by studying the radiolysis of Fricke solution. The study will focus on how dose, dose rate, and initial oxygen concentration influence the activation of the Fenton reaction.

**Methods and Materials:** TOPAS-nBio version 2.0 was used to simulate the radiolysis of the Fricke system. A cubic water phantom of 3µm side was irradiated by 300MeV protons on one of its edges. For UHDR, a proton field (1.5×1.5µm^2^) was delivered in a single pulse of 1ns width. The protons were accumulated until reached 5Gy or 10Gy absorbed dose. For CONV, the independent history approach was used to mimic ^60^Co irradiation. For both dose-rates, oxygen concentrations representative of hypoxic and normoxic tissues (10-250µM) were simulated. The G-value for oxidant ions G(Fe3+) and ΔG-value of Fenton reaction (H_2_O_2_ + Fe^2+^⟶ Fe^3+^+^•^OH+OH^-^) were scored. The simulations ended after G(Fe^3+^) achieved steady-state, and calculated yields were compared with published data.

**Results:** For CONV, G(Fe^3+^) agreed with ICRU-report 34 data by (0.97±0.1) %. For UHDR, G(Fe^3+^) agreed with ICRU data by (1.24±0.1)% and (0.92±0.1)% for 5Gy and 10Gy, respectively. Notably, UHDR at 10 Gy reduced the occurrence of Fenton reactions by (1.0±0.1)% and (11.5±0.1)% at initial oxygen concentrations of 250 µM and 10 µM, respectively. In consequence, UHDR decreased G(Fe3+) by (1.8±0.1)% and (12.5±0.1)% at these oxygen levels. Additionally, increasing the absorbed dose to 15 Gy and 20 Gy at low oxygen (10 µM), UHDR further reduced the ΔG-value by (15.7±0.1)% and (18.6±0.1)%, respectively. The decrease was driven by intertrack effects present in UHDR pulses and its impact on the scavenging effect that oxygen had over hydrogen radicals.

**Conclusions:** UHDR reduces the yield of Fe^3+^ (G(Fe^3+^)) and significantly impacts Fenton reactions, particularly at low oxygen concentrations, while showing minimal effects at higher oxygen levels. This effect becomes more pronounced at higher dose thresholds, such as 10–20 Gy. This emphasizes the important role of the initial oxygen concentration in UHDR and its influence on the Fenton reaction, a mechanism that may contribute to elucidate the FLASH effect.

## 1. Introduction

FLASH radiation therapy (FLASH-RT) has emerged as an innovative approach in the field of radiation oncology. Rediscovered in [1], FLASH-RT utilizes a high dose of radiation, delivered in less than a millisecond, with dose rates exceeding 40 Gy/s [2,3]. This technique has demonstrated promising effectiveness in sparing normal tissues while maintaining comparable tumor control to conventional radiotherapy (CONV-RT), a phenomenon termed the “FLASH effect.” This effect has been demonstrated in multiple preclinical settings, reinforcing the potential of FLASH-RT to transform radiotherapy. Studies on zebrafish embryos, cat and human skin, brain and spinal cord tissues consistently demonstrate the protective effects of FLASH-RT, including reduced tissue damage, faster recovery, and fewer neurological side effects compared to CONV-RT [4–9]. These findings are supported by pioneering clinical trials, including the first FLASH-RT treatment of a patient with subcutaneous T-cell lymphoma, achieving a complete response with minimal toxicities [5]. Similarly, proton FLASH-RT has shown clinical feasibility and comparable safety and efficacy to CONV-RT in the palliative treatment of bone metastases [6], an additional trial currently is being investigated [10].

Despite these promising outcomes, the underlying mechanisms behind the FLASH effect, are not yet fully understood [11]. Hypoxia, often observed in solid tumors because of the poor vascular oxygen supply, is associated with radio-resistance and a poor clinical outcome of tumors [12]. In FLASH-RT context, it was hypothesized that the rapid delivery of UHDR (10^6^ Gy/s) depletes oxygen locally, resulting in a momentary radiation-induced hypoxia due to reoxygenation cannot occur within that time frame [13]. This transient hypoxia limits the accumulation of reactive oxygen species (ROS), reducing DNA damage and protecting normal tissue [14–18]. An additional hypothesis that has generated extensive discussion is the increased rates of free-radical recombination and diffusion induced by high dose-rate [4,9,19,20]. For instance, hydrogen peroxide (H_2_O_2_) is a long-living ROS, that plays a significant role in water radiolysis and contributes to DNA damage e.g., single-strand and double-strand breaks [21]. In pure water, irradiation experiments have shown that dose rate impacts the concentration of (H_2_O_2_) [4,9,22]. However, in a biological system, given the time window during which H_2_O_2_ is produced, the presence of intracellular components plays a crucial role in its main precursor, OH• [23]. In consequence, studies of pure water radiolysis alone are not sufficient to reflect what occurs in biological systems [20].

In simulating biological systems, it is essential to consider other parameters like the intracellular arsenal of antioxidants and the presence of copper and iron metals able to generate OH• from H_2_O_2_, e.g., through the Fenton reaction (oxidation of Fe^2+^ to Fe^3+^ and generation of OH•) [24]. Indeed, as a reservoir of iron ions Fe^2+^, the Labile Iron Pool (LIP) frequently increased in tumor cells compared to normal cells [25] participates in Fenton reactions, and the produced OH• can inflict substantial damage to biomolecules, including DNA, RNA, and proteins [24,26,27]. The oxidized form of guanine, 8-oxo-2′-deoxyguanosine (8-oxo-dG), is usually used as a specific marker of oxidative DNA damage [28]. Interestingly, a positive correlation has been observed between cellular LIP levels and the production of 8-oxod-dG, a marker of ROS-induced DNA damage in human lymphocytes [25,27]. Research from the 1990s showed an inverse dose-rate effect on lipid peroxidation, which was further enhanced with increasing initial concentrations of ferrous ions (Fe^2+^) [29]. More recently [30], used cell-free models to compare lipid peroxidation yields, finding significantly reduced lipid peroxidation under FLASH-RT. Furthermore, combining Dirac pulse dose rates (300 MeV, protons) with cystamine in Fricke solution, reduces G(Fe^3+^) production by scavenging OH• radicals [31], suggesting that combining antioxidants like cystamine with UHDR may enhance the protective effect of FLASH-RT.

While the mechanism driven by LIP may add to the explanation of tumor control with FLASH-RT [13], the differential sparing of normal tissues as compared to CONV-RT requires further investigation. Previous work also demonstrated that the initial oxygen concentration plays a critical role in driving “FLASH effect” [9,30,32]. It is widely suggested that the dose-rate dependence of the Fenton reaction presents a plausible molecular mechanism underlying the FLASH effect [11,33]. Thus, the investigation of the Fenton reactions under FLASH-RT conditions, particularly in relation to varying initial oxygen concentrations, could bring new insights into the underlying mechanisms of the “FLASH effect”.

The Fricke detector is a stable and reliable dosimeter that has a linear response with dose, which makes it a valuable system for dosimetry of ultra-high and low dose-rate irradiation [34–37]. The presence of the Fenton reaction in the Fricke solution makes this system attractive for studying the effects of dose-rate using Monte Carlo track-structure simulations (MCTS). MCTS has been used to model the effect of dose-rate in the production of ROS, allowing the characterization of the spatial and temporal distribution of reactants [31,38,39]. MCTS allows the quantification of chemical yields by modeling the reactions between radiolytic chemical species and their recombination with the bulk, allowing the tracking of direct and indirect actions of radiation with biological targets like DNA [40–44], and the simulation of the reactions occurring in the Fricke solution[31,35,38].

The current study aimed to investigate with MCTS simulations using TOPAS-nBio [38] the effects of UHDR irradiation on Fenton reactions. For that, we propose the use of the Fricke solution to study the implications of these reactions by varying the dose rate and initial oxygen concentration ([O_2_]_0_). The simulations were performed for doses and dose-rates relevant to clinical settings [38]. By exploring the effects of varying [O_2_]_0_ and dose rates in Fenton reactions, the research seeks to contribute to a more comprehensive understanding of the radiochemical mechanisms involved in the “FLASH effect.”

## 2. Methods and materials

Monte Carlo track structure simulations were conducted using TOPAS-nBio version 2.0 [44], built on top of TOPAS version 3.9 [40,45] with Geant4/Geant4-DNA version 10.07. p01 [46–48]. The simulation of water radiolysis with MCTS involves three distinct time stages including physical (10^−15^s), pre-chemical (10^−12^s) and chemical stages (10^−6^s). For the simulation of the Fricke solution, the chemical stage must follow the behavior of chemical species beyond 1µs (non-homogeneous) until a few seconds [44].

It has been shown [38] that for low-LET and low-dose rate conditions, an independent history approximation reproduces the reported chemical yields values, in water, from ICRU [37]. A “history” is defined as the interaction of the primary particle and all its secondary particles with the medium, encompassing the production of all chemical species until the end of the non-homogeneous chemistry stage. The G-value, defined as the number of chemical species produced or lost per 100 eV of energy deposit, can be obtained from the average of multiple independent histories. In this independent history approach, the G value is thus independent of the absorbed dose and dose-rate. On the other hand, for high dose-rate conditions, an accumulated history (from individual histories) approach should be followed. In this approach, the interaction of multiple histories separated in space and time can form a radiation pulse. In this way, an “intertrack effect” in G values can be simulated [38], whereas the accumulation of history tracks until a desired absorbed dose in a given target can be retrieved. The dose-rate can be set by modifying the radiation pulse width. As a result, in this simulation approach, the G value is expected to depend on the absorbed dose and the time duration of the pulse, that is, on the dose rate. For both approaches, the independent reaction time (IRT) method [49,50] can be used to efficiently simulate the homogenous and non-homogenous chemistry stages.

In this work, a cubic water phantom with a volume of 3×3×3 µm^3^ was irradiated with a monoenergetic proton field of 300 MeV (LET 0.3 keV/µm). For UHDR, the initial time of each proton was sampled from a Gaussian distribution to create a radiation pulse of 1 ns FWHM, to mimic an extreme scenario of FLASH-RT conditions and highlight the role of inter-track events. Protons were accumulated until achieve a total dose of 5 Gy and 10 Gy. These dose values are within fractional doses used in stereotactic body radiation therapy, whereas a fractionation of 8 Gy in a single fraction regime was used in the FLASH-01 clinical trial investigating proton FLASH radiotherapy [6,10]. For the independent history approach, each proton history was simulated from time t = 0 s without any time or space overlap between subsequent histories. The protons were distributed over a flat field of 1.5 × 1.5 µm^2^ (i.e., ½ side area of water phantom), incident at zero divergence on one side of the cubic phantom, Figure 1. The target size maintained negligible boundary effects that occurred with smaller target sizes (where radicals diffused away the target), while maintained a good compromise with the computational cost. In addition, a verification tests using a bigger volume of (5 µm)^3^ and a field size of (2.5 µm)^2^ showed minimal differences between the two settings (<0.5%), see Figure1_Ex (Supplementary Data). Negligible effects in chemical yields using a target size of 3 µm side were also confirmed in a separate study [39].

**Figure 1:**
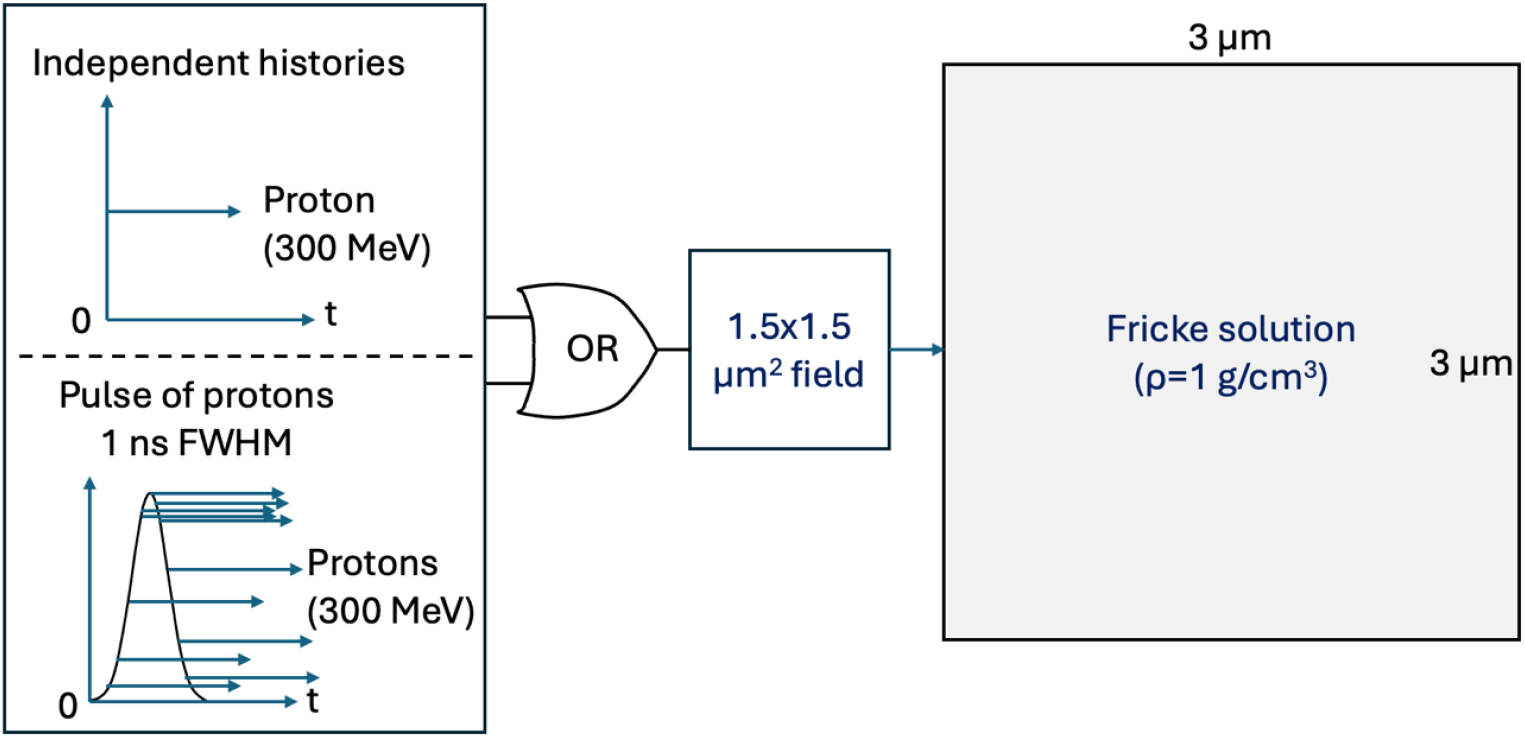
The simulation setup employed in this study. A cubic phantom with 3 µm on each side, serving as the scoring volume filled, was irradiated by monoenergetic protons. The particle beam distribution (zero divergence) within the XY plane spot is set to one half the phantom width. The particles are delivered as independent histories, or in a pulse of 1 ns FWHM.

The physics module TsEmDNAPhysics was used to simulate the track-structure interactions between charged particles and water. This module was derived from Geant4-DNA (v.10.7.p3) constructor and has been extensively described elsewhere [51]. The module includes models for simulating ionization of electrons, protons and ions, electron elastic scattering, vibrational excitation, electron attachment, and thermalization of sub-excited electrons. The TsEmDNAChemistry module was used to activate the radiolysis and chemical process that followed. These modules were validated in TOPAS-nBio for water radiolysis induced by low LET radiation [43]. The key reactions for the Fricke are shown in Table 1, whereas the full list of chemical reactions for water radiolysis and the Fricke solution used in this work were detailed elsewhere [38].

**Table 1:** Key reactions in the Fricke solution responsible for the generation of Fe^3+^, with reaction rates and models adapted from the previous TOPAS-nBio validation [38].

| Main reactions | Reaction rates |
| --- | --- |
| $R1: e_{aq}^- + H^+ \rightarrow H^\bullet$ | $k1 = 2.3 \times 10^{10} \text{ M}^{-1}\text{S}^{-1}$ |
| $R2: H^\bullet + O_2 \rightarrow HO2^\bullet$ | $k2 = 2.1 \times 10^{10} \text{ M}^{-1}\text{S}^{-1}$ |
| $R3: OH^\bullet + Fe^{2+} \rightarrow Fe^{3+} + OH^-$ | $k3 = 3.4 \times 10^8 \text{ M}^{-1}\text{S}^{-1}$ |
| $R4: HO2^\bullet + Fe^{2+} \rightarrow Fe^{3+} + HO2^-$ | $k4 = 7.9 \times 10^5 \text{ M}^{-1}\text{S}^{-1}$ |
| $R5: H2O2 + Fe^{2+} \rightarrow H + Fe^{3+} + OH^- + OH^\bullet$ | $k5 = 52 \text{ M}^{-1}\text{S}^{-1}$ |
| $R6: SO4^- + Fe^{2+} \rightarrow Fe^{3+} + SO4^{2-}$ | $K6 = 2.79 \times 10^8 \text{ M}^{-1}\text{S}^{-1}$ |
| $R7: H^\bullet + SO4^- \rightarrow HSO4^-$ | $K7 = 1 \times 10^{10} \text{ M}^{-1}\text{S}^{-1}$ |
| $R8: H^\bullet + S2O8^{2-} \rightarrow SO4^- + HSO4^-$ | $K8 = 2.5 \times 10^7 \text{ M}^{-1}\text{S}^{-1}$ |
| $R9: OH^- + HSO4^- \rightarrow SO4^-$ | $K9 = 1.5 \times 10^5 \text{ M}^{-1}\text{S}^{-1}$ |
| $R10: e_{aq}^{-1} + S2O8^{2-} \rightarrow SO4^- + SO4$ | $K10 = 1.2 \times 10^{10} \text{ M}^{-1}\text{S}^{-1}$ |
| $R11: H2O2 + SO4^- \rightarrow HO2^\bullet + SO4^{2-}$ | $K11 = 1.2 \times 10^7 \text{ M}^{-1}\text{S}^{-1}$ |
| $R12: OH^- + SO4^- \rightarrow OH^\bullet + SO4^{2-}$ | $K12 = 8.3 \times 10^7 \text{ M}^{-1}\text{S}^{-1}$ |
| $R13: SO4^- + SO4^- \rightarrow S2O8^{2-}$ | $K13 = 4.4 \times 10^8 \text{ M}^{-1}\text{S}^{-1}$ |

The simulation parameters for the aerated Fricke solution consisted of 1 mM ferrous (Fe^2+^) ions and 0.4 M sulfuric acid (H_2_SO_4_, pH = 0.46). The initial oxygen concentration ([O_2_]_0_) was varied between 10 and 250 µM, concentrations representative of hypoxic and normoxic tissues, to explore its influence on the Fenton reaction (R5) in Table 1. The time evolution of G-values for Fe^3+^ (G(Fe^3+^)) and ΔG-values were reported. The ΔG-value was defined as the number of reactions N_R_ of a kind that occurred per 100 eV of energy deposit:

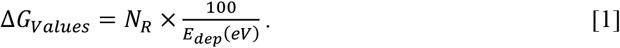

All the quantities scored from the Monte Carlo simulations had 0.1% of statistical uncertainty and presented at 1 standard deviation. The differences between independent histories and UHDR were calculated as:

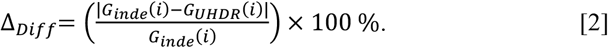

## 3. Results

### 3.1. Model Verification: Independent Histories and UHDR Approach

An initial consistency checks of pulse delivery and absorbed dose was conducted by comparing independent tracks and UHDR for a 1 Gy dose. ICRU report indicates that a dose of 1 Gy per pulse at UHDR does not impact the Fe^3+^ yield compared to a low dose rate irradiation [37]. Our simulation setup showed differences below 0.5%, at steady state, see Figure2_Ex (supplementary data), thereby confirming that the independent history method mimics low or conventional dose rate scenario. Later, we compared the TOPAS-nBio simulated G(Fe^3+^) yields with ICRU Reports [37] for the aerated Fricke solution ([O_2_]_0_ = 250 µM) under conventional (independent histories) and UHDR irradiation (5-10 Gy per pulse).

As shown in Figure 2, the simulated G(Fe^3+^) agreed with ICRU report data [37] by 0.97% for independent histories and 1.24% and 0.92% for pulsed irradiation at 5 Gy and 10 Gy per pulse, respectively. On the other hand, the deaerated Fricke solution was compared with previously published data [31] for only independent histories due to the lack of measured data at such low oxygen concentration. The comparison with data from [52], showed a difference of 4.36% at 200s, as depicted in the blue (right size, Figure 2). The time evolution of G(Fe^3+^) for the aerated and deaerated Fricke models from 1 ps to 200s, from this work and from published MC data [31] are shown in Figure 3. In the figure, results of measured data for aerated and deaerated are also shown *[37,52]*. As depicted, the MC simulations behave similarly along the entire period of time.

**Figure 2:**
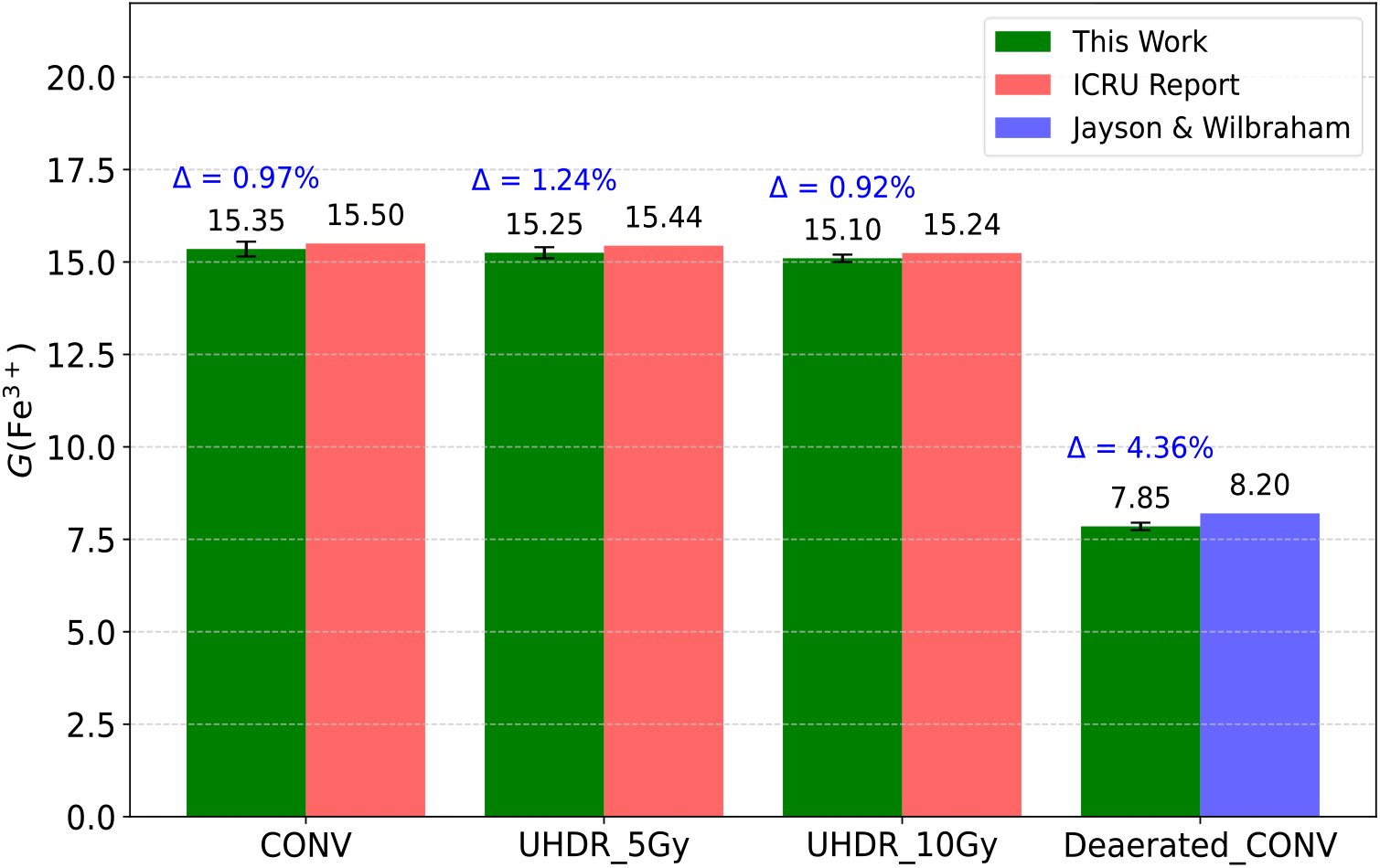
G(Fe^3+^) calculated in this work (green bars) and the ICRU report data [37](red bars), for CONV, and UHDR irradiations at 5 Gy and 10 Gy for 250 µM [O2]_0._ For deaerated Fricke solution (0 µM [O2]_0_), G(Fe^3+^) calculated in this work (green bar) and from [52] (blue bar) are shown in the right size of the figure. The uncertainties for each scenario are represented by error bars, with percentage differences (Δ) between literature and this work displayed above the bars

**Figure 3:**
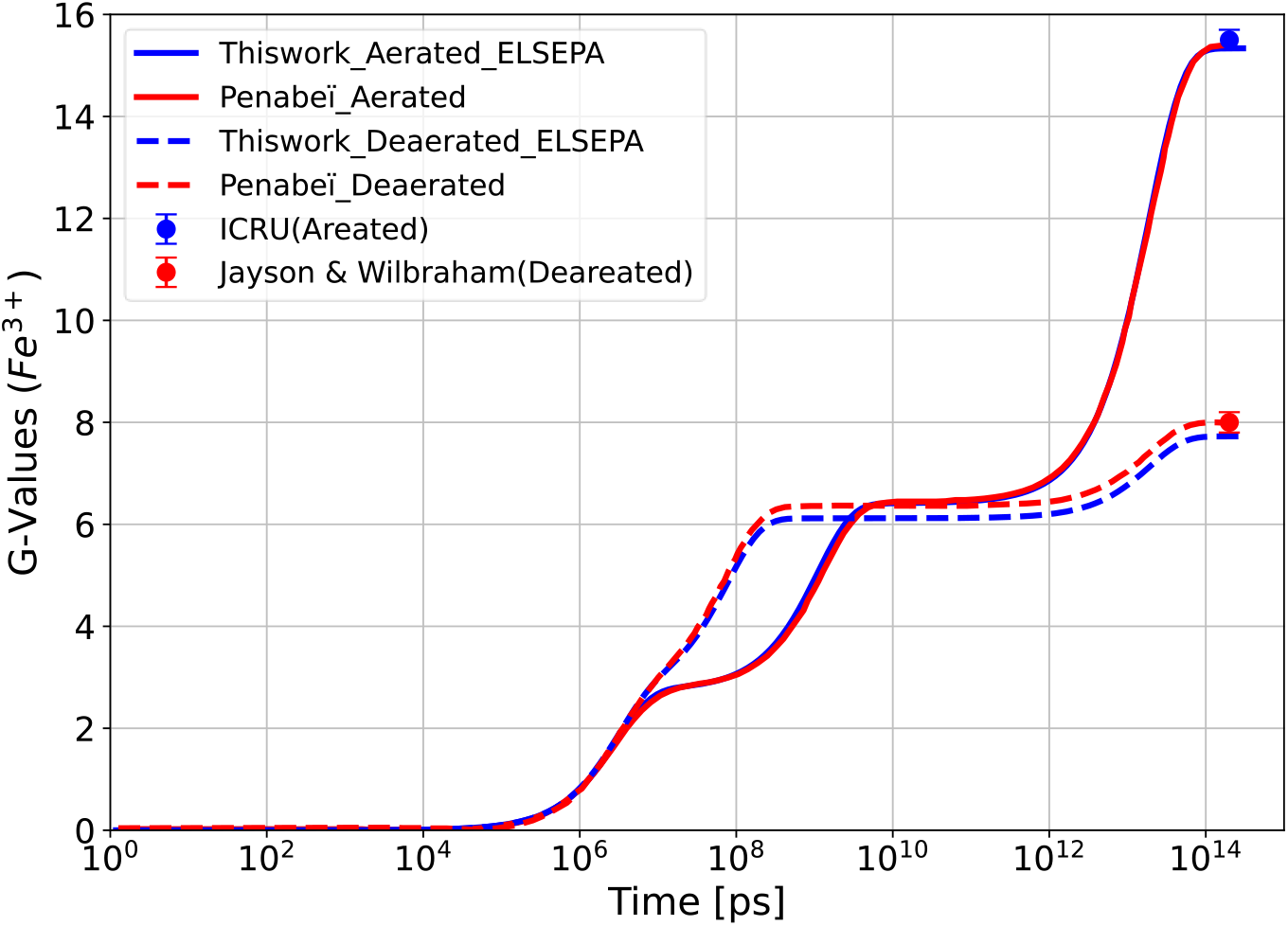
Time evolution of G(Fe^3+^) at 250 µM [O_2_]_0_ for independent histories from this work (blue solid lines) and Monte Carlo data from [31] (red solid line). The ICRU-reported value of 15.5 ± 0.2 Fe^3+^/100 eV for 250 µM is indicated by the blue solid symbol [37] at 200 s time point (i.e., steady state). Additionally, a comparison is shown for G(Fe^3+^) at 0 µM [O_2_]_0_ between Monte Carlo data from [31] and this work models (ELSEPA), along with data from [52] represented by a red solid symbol at steady state.

### 3.2. Effect of UHDR on G-value of Fe^3+^

The time evolution of the G(Fe^3+^) for both UHDR at 10 Gy (red lines) and independent histories (blue lines) at two [O_2_]_0_ (10 µM and 250 µM) are shown in the left and right panels of Figure 4, respectively. Point-to-point differences computed with Equation 2 are shown in the bottom of each panel. As shown, at 10 µM of [O_2_]_0_ UHDR irradiation reduced G (Fe^3+^) by 12.5±0.1% (at 10^14^ps). At 250 µM of [O_2_]_0_, UHDR irradiation reduced by 1.8±0.1 % at (10^14^ ps) but it had a slightly higher reduction ∼5%, at 10^7^ ps.

**Figure 4:**
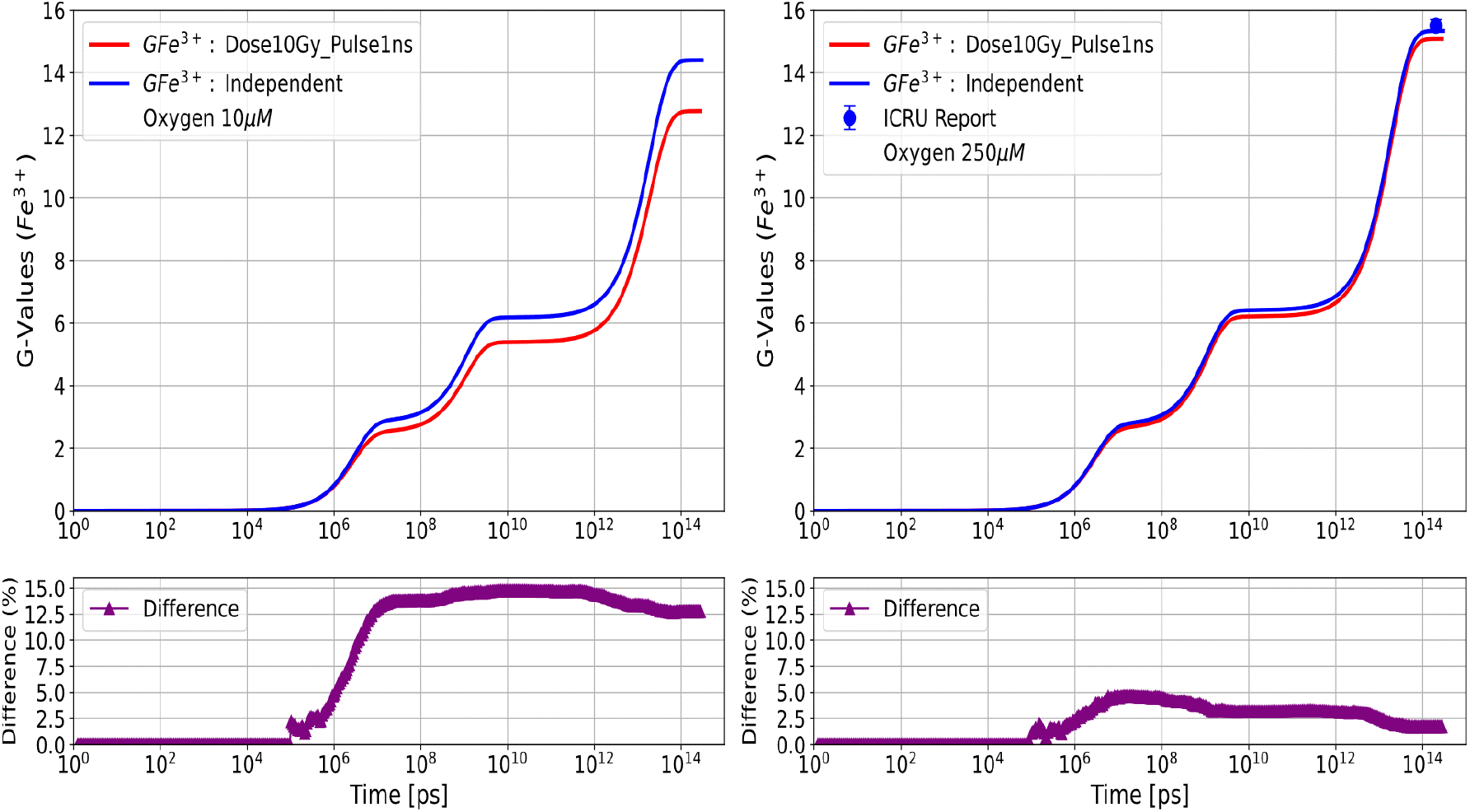
Temporal evolution of G(Fe3+) calculated with independent histories (blue curves), and UHDR for 10 Gy of absorbed dose (red curves). Left panel, G(Fe^3+^) for an initial oxygen concentration of 10 µM. Right panel: G(Fe^3+^) for an initial oxygen concentration of 250 µM, the ICRU-reported value for 250 µM (15.5±2) is shown with the solid symbol [37]. The bottom of both panels shows point-to-point percentage differences.

To explore the main reactions contributing to the decrease of G(Fe^3+^), the ΔG-values for those reactions with the greater contribution to the Fricke system are presented in Figure 5. Contributions from other reactions were less than 1% across the entire time range (10^6^ ps to 10^14^ ps). In general, at [O_2_]_0_ of 10 µM, the effect of UHDR at 10 Gy on the time evolution of ΔG-values is more pronounced compared to 250 µM throughout the entire time range, reaching the largest difference in ΔG-values of 15.5%. More precisely, the reduction in G(Fe^3+^) at low [O_2_] _0_ is primarily driven by reactions R2 by Δ_*Diff*_ up to 15.5 % and R3 (11.5%) starting from 10^6^ ps and sustained by reactions R4 (15.5%) and R5 (11.5%) at later time points 300 s, this sequence is depicted in Figure 5.

**Figure 5:**
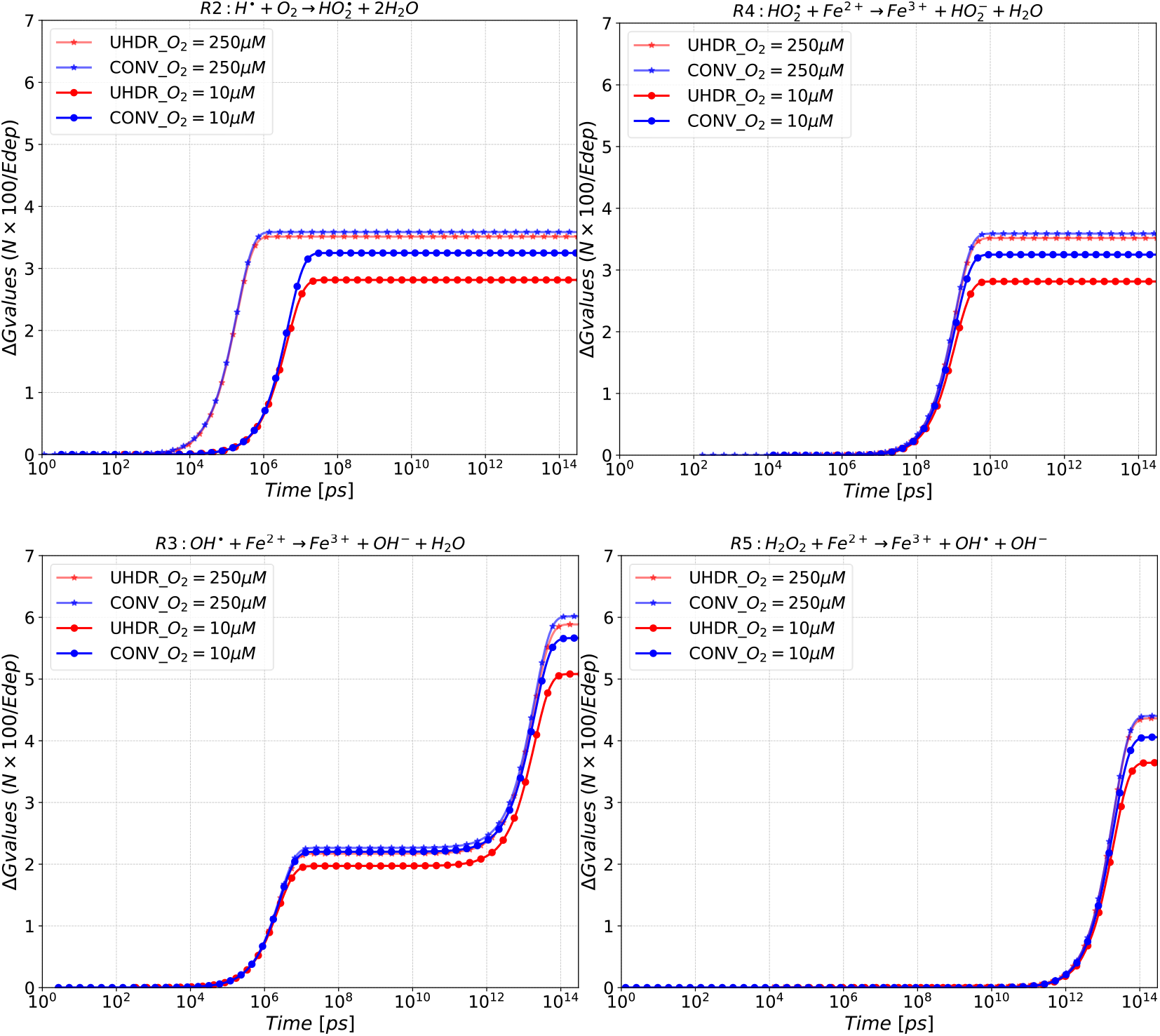
Temporal evolution of ΔG-values for reactions that contribute mostly to the decrease in G(Fe^3+^). The data compares the effects of UHDR at 10 Gy (red lines) and Independent Histories (blue lines) under two oxygen tension conditions at 10 µM (circular marker) and 250 µM (star marker).

### 3.3. Impact of Initial Oxygen Concentration and UHDR on Fenton Reactions

The impact of UHDR at 5 Gy and 10 Gy in G(Fe^3+^) at steady state, quantified by Equation 2, as a function of the initial oxygen concentration (10-250 µM) is shown in Figure 6. As depicted, the impact of UHDR decreased monotonically with increased oxygen concentration, from a maximum difference of 6.7 ± 0.1% and 12.5 ± 0.1% under low oxygen conditions ([O_2_]_0_ = 10 µM) to 1.3 ± 0.1% and 1.8% at high oxygen conditions ([O_2_]_0_ = 250 µM), for 5 Gy and 10 Gy, respectively.

**Figure 6:**
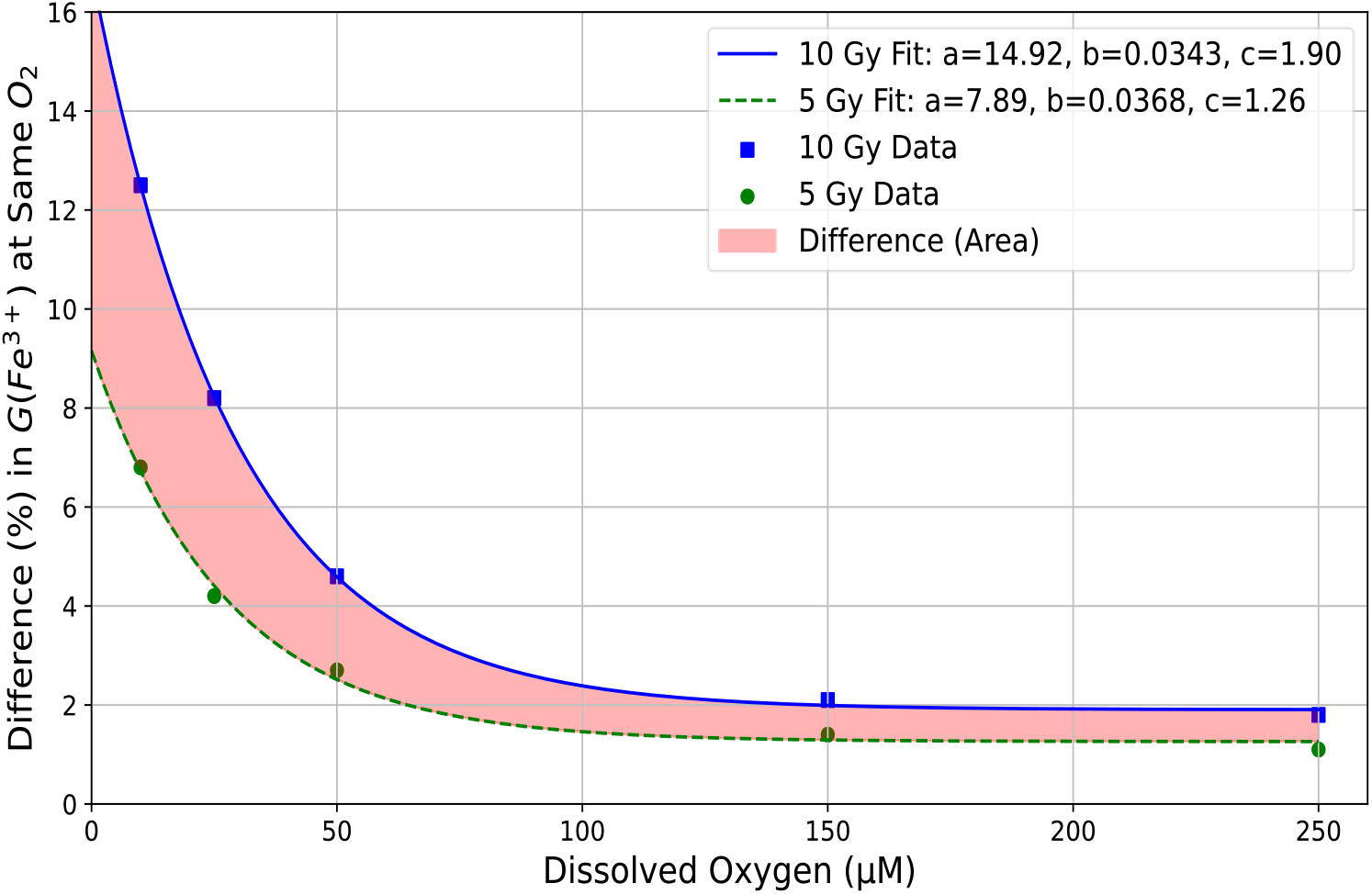
G(Fe^3+^) difference (%) between UHDR and CONV as a function of initial oxygen. The blue solid line and squares represent the 10 Gy dose, indicating the fitting function and data points, respectively. The green dashed line and circles represent the 5 Gy dose, showing the fitting function and data points, respectively. The difference driven by the total doses (5 to 10 Gy) is depicted by the colored area.

Furthermore, the G(Fe^3+^) difference (%) as function of [O_2_]_0_ followed an exponential law decay as a function of [O_2_]_0_. The reduction rate was obtained from a fit using *Rate* = −*a e*^−*b*⋅[*o*^_2_^]^_0_, where a is the initial difference at low oxygen [O_2_]_0_, and b is the rate. From that fit, for 10 Gy at [O_2_]_0_ =50 µM, the reduction rate around Rate =−0.09 and reduced to −0.046 for 5Gy, details on the calculation, see Section4 in Supplementary Data.

The effect of UHDR at 10 Gy on the reactions that contribute most to G(Fe^3+^) including Fenton reaction (quantified with the ΔG-values difference), as a function of oxygen concentration is shown in Figure 7. As depicted, the differences in ΔG-values decreased as [O_2_]_0_ increased. For instance, the Fenton reaction (R5) occurred 11.5% less frequently at UHDR conditions compared to CONV at 10 µM [O_2_]; and became comparable (around 1 %) at 250 µM [O_2_]_0_. Notably, R2 drove the reactions with oxygen, which depends on its availability. Under UHDR conditions, oxygen consumption was reduced by more than 15.5% compared to CONV-RT at low oxygen levels. However, at 250 µM [O_2_]_0_, the higher competition between oxygen and intertrack events for H•, led to a difference around 2%. On other hand, Figure 7 shows that R2 and R4 overlap, as the products of R2 act as the reactants for R4, driving the formation of H_2_O_2_ required for R5. This sequence is confirmed by the similar reduction rates of R4 and R5 as a function of oxygen.

**Figure 7:**
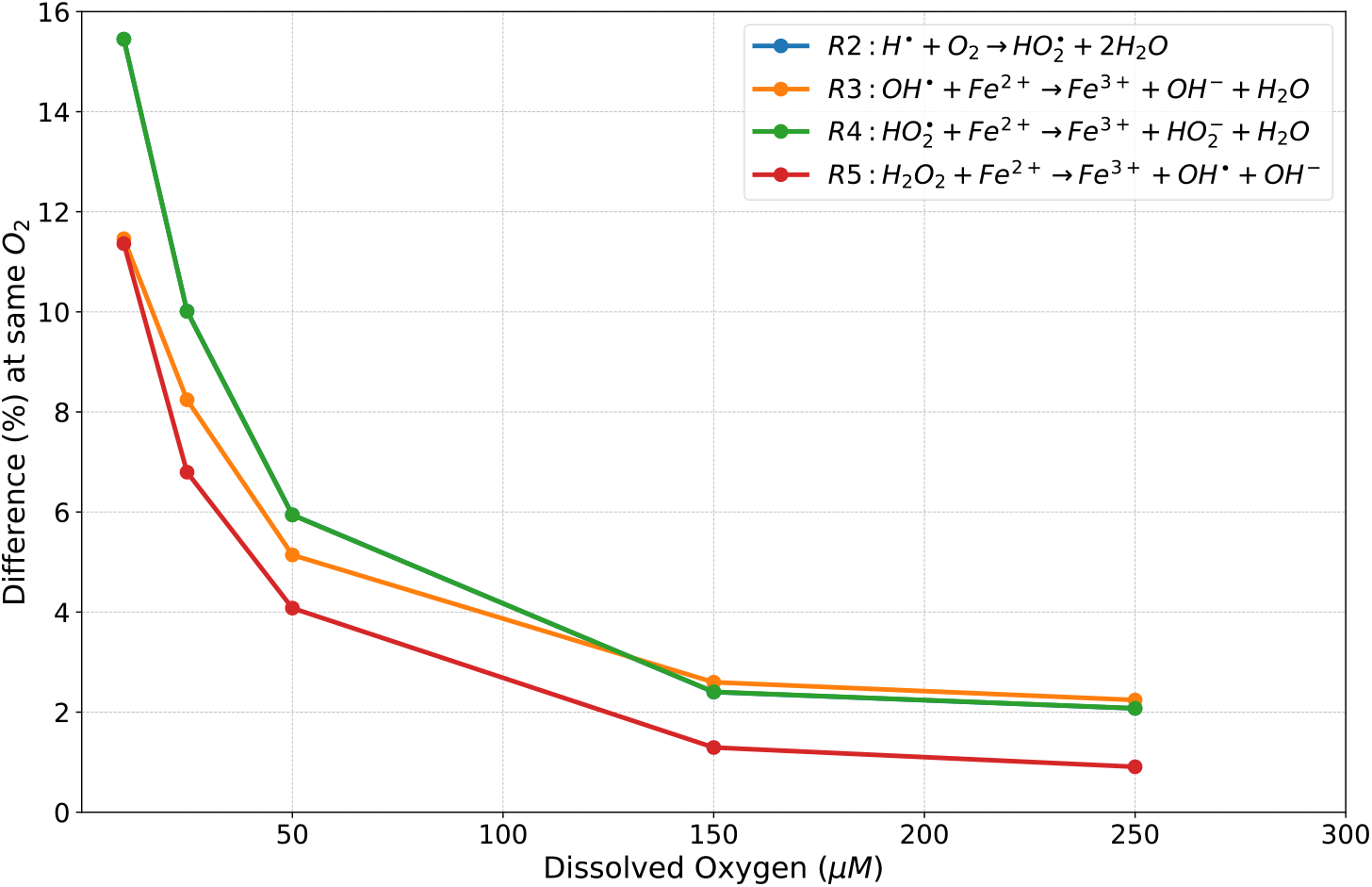
This figure quantifies the difference in ΔG-values of reactions under Ultra-High Dose Rate and Independent histories approach across a range of initial oxygen concentrations [10-250 µM], R2 and R4 overlapped.

These findings show that the impact of UHDR on the reduction of reactive species and Fenton reactions becomes more pronounced at lower oxygen levels. The significance of these results will be discussed in detail, with a focus on the mechanistic differences in oxygen consumption and radical dynamics under UHDR compared to independent histories.

## 4. Discussion

This study used Monte Carlo track-structure simulations to investigate the correlation between UHDR irradiation and initial oxygen concentration ([O_2_]_0_) on the Fenton reaction through the Fricke solution, with a 300 MeV proton beam under UHDR and CONV-RT. Clinically relevant absorbed doses of 5 Gy and 10 Gy were used to quantify the G(Fe^3+^) yields and ΔG-values, employing the independent history approach for CONV-RT and a pulsed beam for UHDR at varying [O_2_]_0_. The findings of this study demonstrate a clear dependence of G(Fe^3+^) yields on [O_2_]_0_ under UHDR compared to CONV-RT. Specifically, the G(Fe^3+^) yield exhibited a maximum decrease of 12.5% ± 0.1% at [O_2_]_0_ = 10 µM, with a minimal reduction of 1.8% ± 0.1% at [O_2_]_0_ = 250 µM under UHDR relative to CONV-RT.

Tests were performed to verify the consistency of the simulation setup. We selected a target size that resulted in negligible border effects in G values, and that was previously used in separate studies [39,53]. Whenever available, comparisons with measured data were performed. The simulated yield for the aerated Fricke at a low dose rate was within 1% from the data reported by ICRU [37], and the use of low doses (1 Gy) yielded results equivalent to the independent history approach. For deaerated Fricke solution, the TOPAS-nBio G(Fe^3+^) data was 4.3% lower than the measured data from [52]. This difference was in part attributed to the chosen electron elastic scattering model. For instance, we performed simulations using the so-called Champion elastic scattering model available in Geant4 (based on the partial wave theory) [54], leading to a difference of 2.5% for the deaerated Fricke solution. Nevertheless, all our simulation results used ELSEPA model for elastic scattering, which considers relativistic corrections and is currently recommended by the Geant4-DNA and TOPAS-nBio collaborations [43,55].

Most solid tumors are hypoxic [56], with oxygen concentrations as low as 1% [12]. For our study, we selected oxygen concentrations that correspond to e.g., 1%, 5% and 20% for 10 µM, 50 µM and 250 µM, respectively. These values correspond to physiological levels of oxygen found in tumors (∼1%) and healthy tissues (5%-20%). While our study was performed in an aqueous solution rather than a cellular environment, our simulation setup provides a relevant framework for analyzing the relative impact of oxygen and dose-rate in the Fenton reaction at similar oxygenation levels than tumor and normal cell environments. The dynamic interplay between dose rate, doses (5-10Gy) and oxygen availability shown in Figure 6, demonstrates that the impact of UHDR on G(Fe^3+^) yields diminishes as the initial oxygen increases (10-250 µM), emphasizing the dependence on oxygen concentration. However, our model did not consider the reoxygenation process that indeed exists in a biological system.

The contribution of intertrack effects was explored with the ΔG-values for the primary reactions contributing to this observed effect. At higher oxygen concentration (250 µM), the low decrease in G(Fe^3+^) was attributed mainly to the reduction of OH• radicals yield caused by inter-track effects. Specifically, the occurrence of reaction R3 decreases by 2%, while other reactions remain indistinguishable under both modalities (i.e., the difference <1 %), as depicted in Figure 5. The inter-track effect, caused by the overlap of subsequent proton histories delayed in space and time, increased by UHDR at the early chemistry stage, leading to reduction in G(OH•) by 6% at 10^6^ ps. This value aligns with previously calculated yields presented elsewhere [19,22,38].

At low oxygen concentration ([O_2_]_0_ =10 µM), the reduction in G(Fe^3+^) by 12.5% was primarily driven by reactions R2 and R3, starting from 10^6^ ps and sustained by reactions R4 and R5 at later time points in the millisecond range (Figure 5). This reduction was further influenced by the diminished generation of OH• during the early stage (Spur theory), with intertrack effects contributing up to a 14.5% difference, impacting reaction R3. These findings align with previous findings reported in [38]. Furthermore, reaction R2 occurred in lower frequency under UHDR conditions at [O_2_]_0_ =10 µM, compared to the independent histories approach [18,57,58]. This was driven by intertrack effects that reduced hydrogen radicals (H•) at the heterogenous chemistry stage, leading to negligible competition with O_2_ given the low concentration (low scavenging capacity) and thus decreasing the production of HO_2_. As a result, a 25% difference at 10^6^ps was observed. The impact of high dose-rate in hydrogen was also reported in [59]. Consequently, reduced HO_2_ resulted in a subsequent reduction in H_2_O_2_ accumulation under UHDR compared to CONV, see Figure3_Ex and Figure4_Ex (Supplementary data). These combined behaviors resulted in a reduced activation of the Fenton reaction. Therefore, a low occurrence of Fenton reactions driven by low oxygen and UHDR might lead to a reduced generation of OH• at later times, which are a highly reactive radical that drives the indirect damage to DNA [24].

At 10 µM oxygen concentrations, the ΔG-value of the Fenton reaction (R5) between UHDR and CONV reduced monotonically with increasing the dose. For the range of doses 1 Gy to 20 Gy a non-linear relationship was found, as shown in Figure 8. We expect that our model reflects pronounced effects at higher doses and also at different configurations of pulses. Nevertheless, given the high computational cost at higher doses and multiple pulses, these scenarios were not considered but are being studied in a separate work using the Gillespie algorithm, which is still not publicly available in TOPAS-nBio.

**Figure 8:**
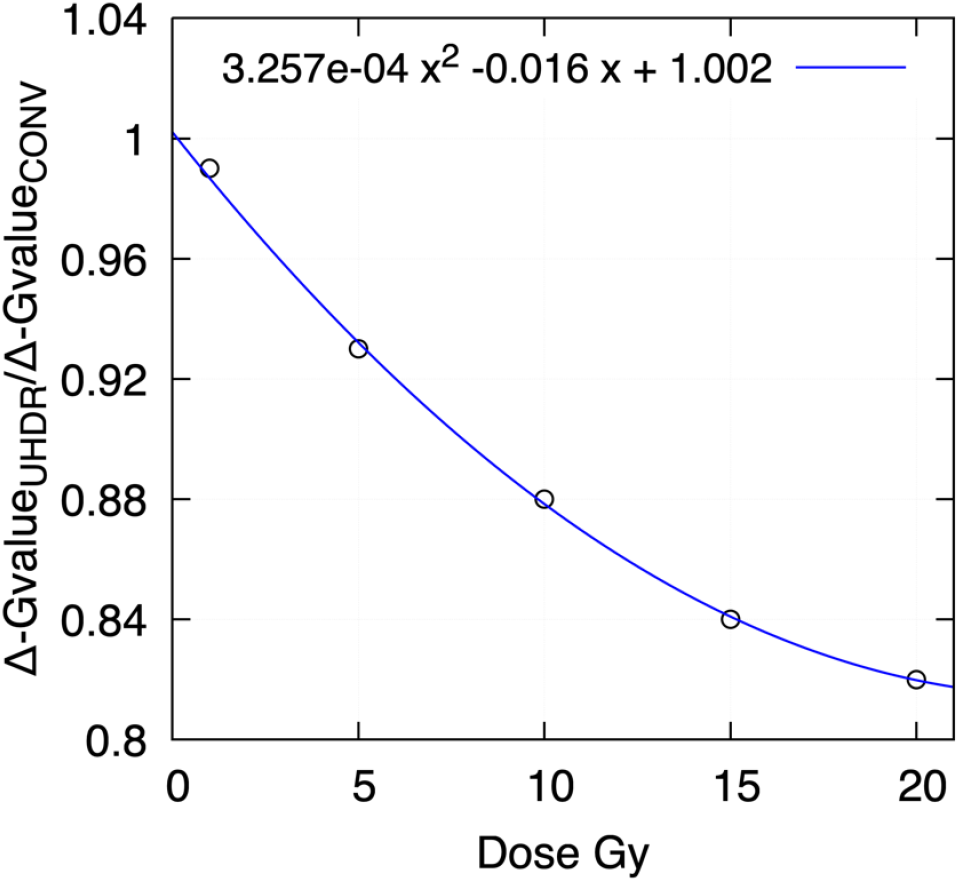
Ratio between ΔG-value at UHDR to ΔG-value at CONV as a function of the dose for 10 µM of oxygen concentration.

An important remark with the current work is that, while Fenton reactions occur in biological systems, our focus is on the dose rate effects studied in the Fricke solution, which also relies on the Fenton reaction. We recognize the distinctions between these two contexts and cautioned the extrapolation of current quantifications with those from biological system.

In this work, high-LET radiation was not considered. The LET value of 0.3 keV/µm used in this work played in favor of producing higher yields of chemical species and the prolonged occurrence of inter-track effects. For example, the G value of hydrated electrons is about 5 times higher for 60-Co than for carbon-ions of 30 MeV at 10^9^ s^-1^ scavenging capacity (about a cellular-like scavenging capacity) [62]. And the inter-track interactions of high-LET ions last only for the duration of the short radiation pulse whereas for low-LET the inter-track spans several orders of magnitude in time [42].

We expect that our findings motivate the experimental quantification of the Fenton reaction under FLASH-RT conditions in a biological system as recently performed for radiobiologically relevant scavengers [31]. Compared to the work in [31], we used absorbed doses commonly used in clinical scenarios for stereotactic body radiation therapy and included the time delay of subsequent proton tracks to produce a pulse of a given width. Nevertheless, more computational development efforts are required to simulate higher absorbed doses, and multiple pulse configurations within long time scales which increase the computation costs and posed a significant limitation in this work.

## Conclusion

In this study, we examined the impact of UHDR irradiation and initial oxygen concentrations on the Fenton reaction. A significant decrease in ferric ion (Fe^3+^) yields was computed of 12.5% at low oxygen levels and less than 1.8% at high oxygen levels, under UHDR compared to CONV dose rates. This reduction was primarily attributed to a decrease in the activation of the Fenton reaction by the reaction between hydrogen atoms and oxygen with a modest contribution of intertrack effects, resulted in 11.5% decrease at low oxygen and less than 2% at high oxygen levels. This highlights the critical role of initial oxygen concentration in FLASH radiotherapy and its influence on the Fenton reaction, a mechanism that may help explain the FLASH effect.

## Acknowledgments

This research was supported through computational resources of HPC-MARWAN (hpc.marwan.ma) provided by the National Center for Scientific and Technical Research (CNRST), Rabat, Morocco.

The authors would like to extend their gratitude to Professor Rabii Ameziane El Hassani for engaging in insightful discussions and sharing valuable ideas, which have contributed to enhancing the quality of this paper.

