## Supplementary data for "Monte Carlo track-structure simulation of the impact of Ultra-Hight Dose Rate and oxygen concentration on the Fenton reaction"

This supplementary data to support the finding in original draft "Impact of Ultra-Hight Dose Rate and oxygen concentration on Fenton reaction: a TOPAS-nBio study"

### 1 Model comparison and validating setup:

Validation tests with an increased volume of (5 µm)³ and a beam size of (2.5 µm)², to confirm low boundary effects where radicals could diffuse. Interestingly, results for 10 Gy at 10 µm oxygen, showing minimal differences (0.5%) compared to (3µm)³ volume  across all time points (1 ps to 300 s), see Figure1_Ex. Furthermore, the aerated Fricke model was validated under UHDR (i.e., at 1 Gy per pulse) conditions, as ICRU reports indicate that a total dose of 1 Gy per pulse does not impact the $Fe^{3+}$ yield compared to independent histories [1]. The differences, shown in the bottom panel *Figure2_Ex*, remain consistently below 0.5%, demonstrating the model's stability and reliability for low dose rates and high dose rates.

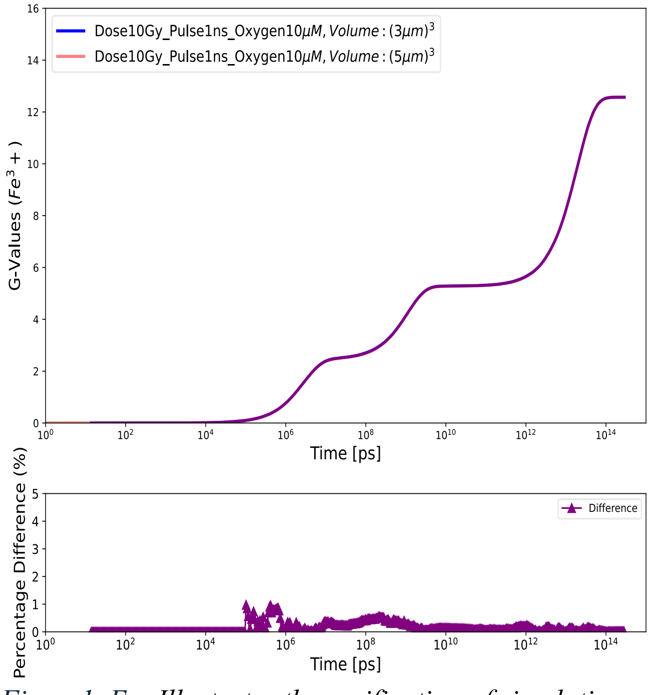


*Figure1_Ex: Illustrates the verification of simulation results under varying beam sizes and volumes. The analysis compares the default setup of a cubic volume of $(5\ \mu m)^3$ with a beam size of $(2.5\ \mu m)^2$, and a smaller setup of a cubic volume of $(3\ \mu m)^3$ With a beam size of $(1.5\ \mu m)^2$. For 10 Gy with 10 µm oxygen, differences below 0.5% at steady state, demonstrating no boundary effects for the smaller target size.*

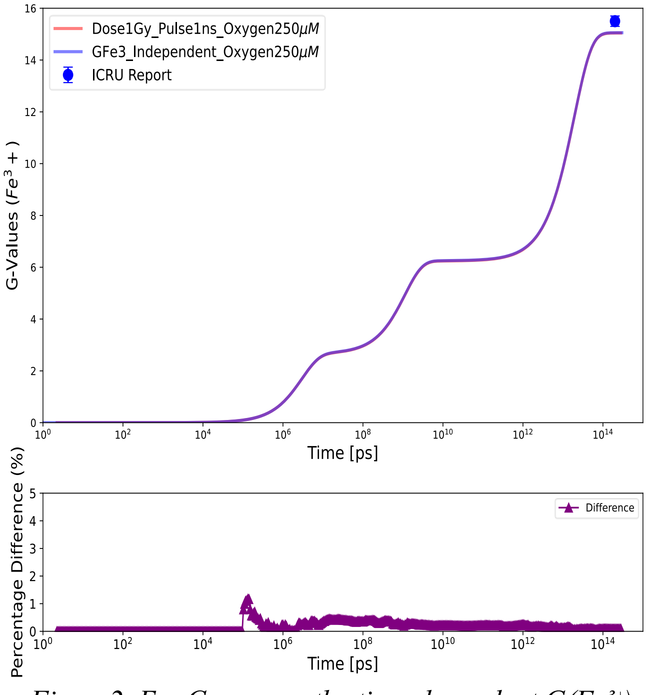


*Figure2_Ex: Compares the time-dependent $G(Fe^{3+})$ yields under UHDR (red line, 1Gy per pulse) conditions and independent histories (blue line) from 1 ps to 300 s. The differences, shown in the bottom panel, remain consistently below 0.5%, the ICRU-reported value 15.5±0.2 $Fe^{3+}$/100eV is shown with the solid symbol.*

### 2 Chemical yield evolution and ΔG-Values:

To dive into the observed decrease under UHDR (10 Gy, pulse 1ns), we report G-Value of the main species contributing to the observed decrease from the early stage $10^6$ps, see *Figure3_Ex*.

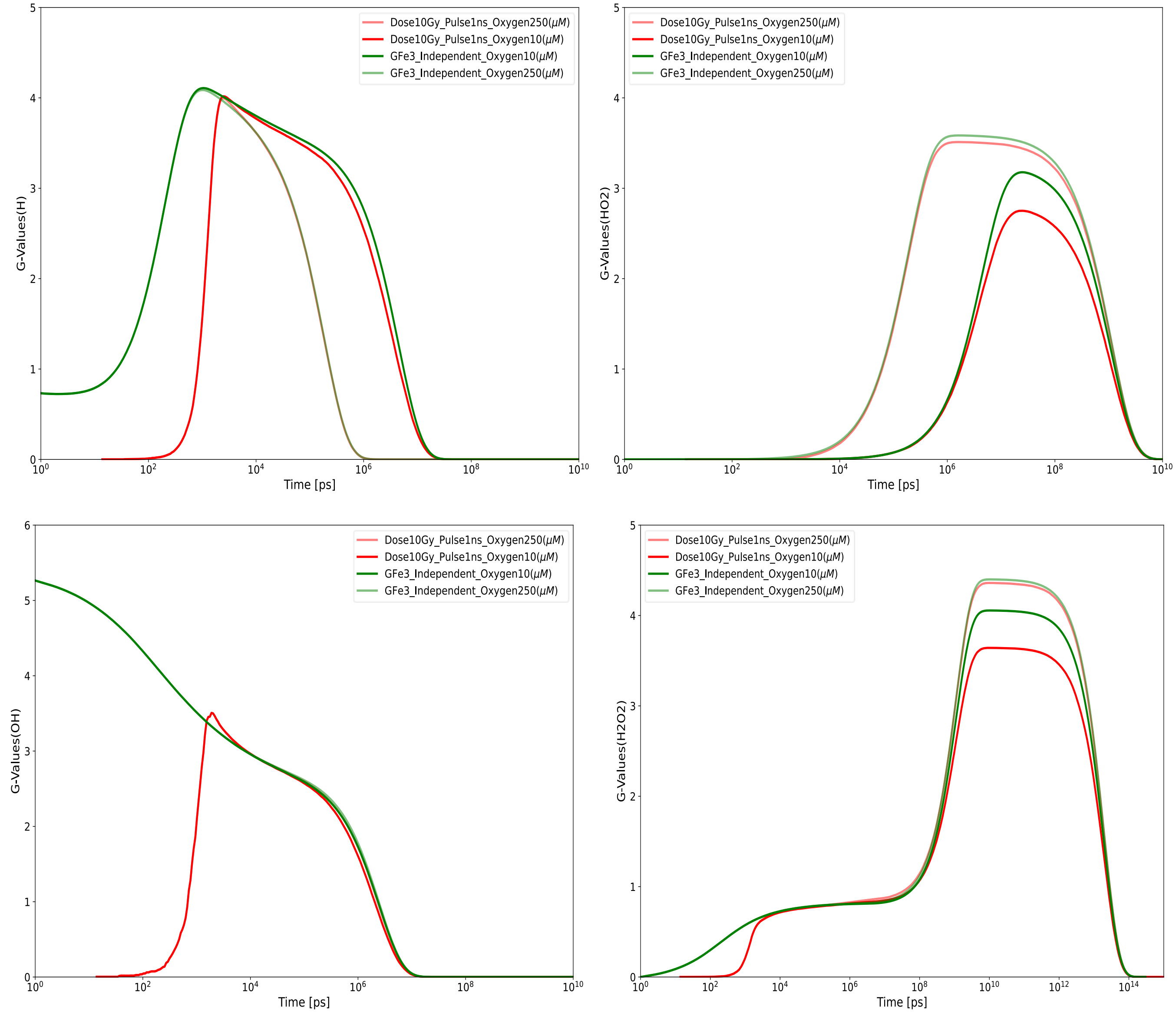


*Figure3_Ex: Chemical species (H2O2, HO2, H, and OH) G-value as function of time for UHDR (10 Gy) red graphs and Independent Histories green, both for normoxic oxygen 20% (250 µM, light opacity) and Hypoxia 1 % (10 µM, high opacity).*

Beside the ΔG-Values of reactions sustain these differences up to late time scale $10^{12}$ps (R3: OH• + $Fe^{2+}$ ⟶ $Fe^{3+}$ + $OH^{-}$) and producing OH• (R5: $H_2O_2$ + $Fe^{2+}$ ⟶ $H^{+}$ + $Fe^{3+}$ + $OH^{-}$ + OH•) as a function of time *Figure4_Ex*.

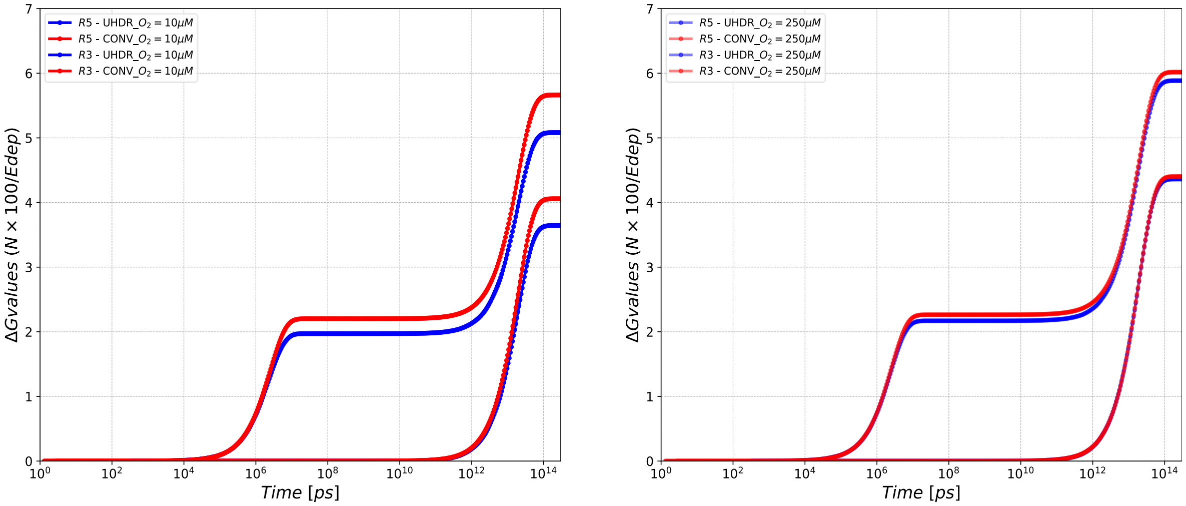


*Figure4_Ex: Illustrates the ΔG-Values of Fenton reactions (R5: $H_2O_2$ + $Fe^{2+}$ ⟶ $H^{+}$ + $Fe^{3+}$ + $OH^{-}$ + OH•) and the scavenging of OH• (R3: OH• + $Fe^{2+}$ ⟶ $Fe^{3+}$ + $OH^{-}$) as a function of time. The left panel, under hypoxic conditions (10 µM), UHDR (blue graphs) compared to CONV-RT (red graphs) at total dose of 10*

### 3 Total dose effect and reduction rates difference (%) of G(Fe3+)

Fitting the G($Fe^{3+}$) difference (%) data points at steady state for 5Gy and 10Gy at initial $[O_2]_0$ in the range [10-250 µM], an exponential decay function is extracted:

$$Difference = a.exp(-b.[O_2]_0) + c.$$

Where parameter b governs how quickly the difference decays with increasing initial oxygen$[O_2]_0$, and its value directly influences the reduction Rate. a: Represents the initial difference at low oxygen concentrations. **c**: The baseline value, which represents asymptotic behavior at very high oxygen concentrations.

Differentiating the exponential decay Function one has:

$$Rate = -a.b.exp(-b.[O_2]_0).$$

This will give the rate of change of the difference between UHDR and independent histories at the specific oxygen concentration.

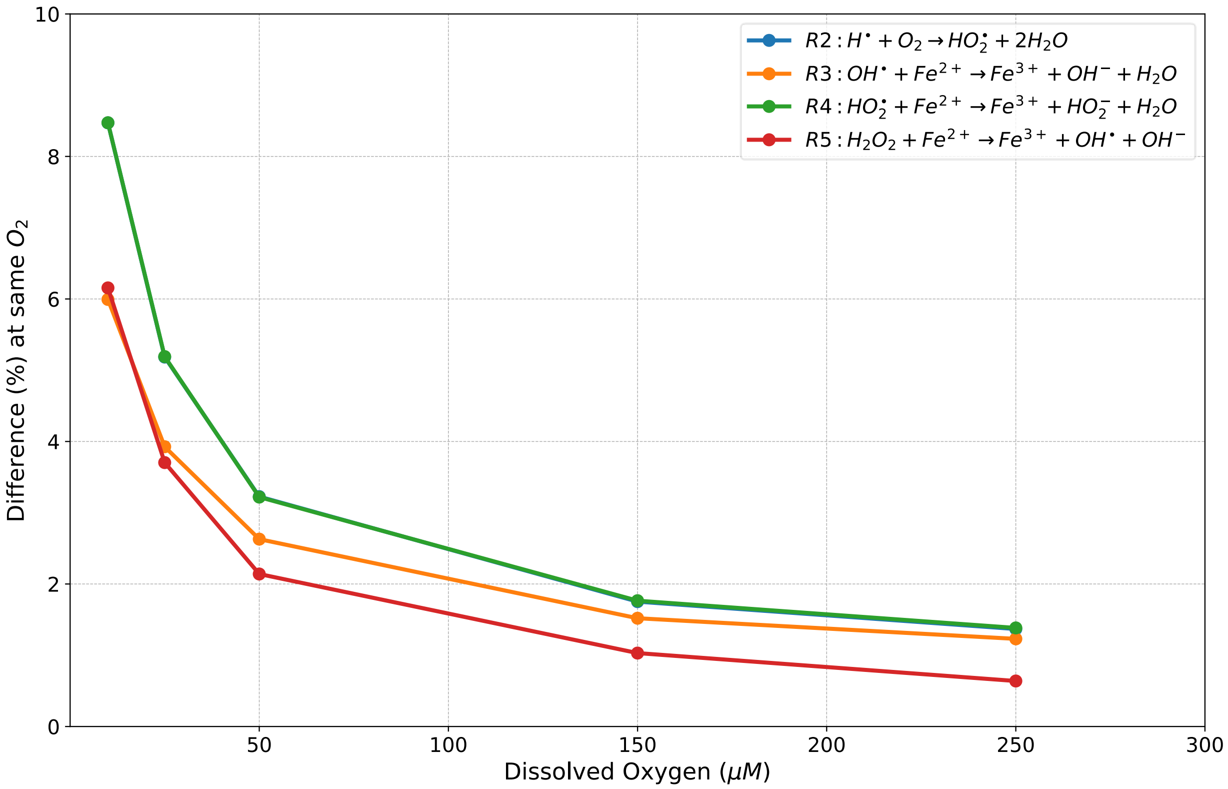


*Figure5_Ex: This figure illustrates the behavior of the Fenton reactions under UHDR conditions with a total dose of 5Gy. Wherein R2 and R4 data point are overlapped since R2 yields initialize R4.*
